# Development of an in vitro model for animal species susceptibility to SARS-CoV-2 replication based on expression of ACE2 and TMPRSS2 in avian cells

**DOI:** 10.1101/2021.08.18.456916

**Authors:** Darrell R. Kapczynski, Ryan Sweeney, David L. Suarez, Erica Spackman, Mary Pantin-Jackwood

## Abstract

The SARS-CoV-2 (SC2) virus has caused a worldwide pandemic because of the virus’s ability to transmit efficiently human-to-human. A key determinant of infection is the attachment of the viral spike protein to the host receptor angiotensin-converting enzyme 2 (ACE2). Because of the presumed zoonotic origin of SC2, there is no practical way to assess every species susceptibility to SC2 by direct challenge studies. In an effort to have a better predictive model of animal host susceptibility to SC2, we expressed the ACE2 and/or transmembrane serine protease 2 (TMPRSS2) genes from humans and other animal species in the avian fibroblast cell line, DF1, that is not permissive to infection. We demonstrated that expression of both human ACE2 and TMPRSS2 genes is necessary to support SC2 infection and replication in DF1 and a non-permissive sub-lineage of MDCK cells. Titers of SC2 in these cell lines were comparable to those observed in control Vero cells. To further test the model, we developed seven additional transgenic cell lines expressing the ACE2 and TMPRSS2 derived from *Felis* (cat), *Equus* (horse), *Sus* (pig), *Capra* (goat), *Mesocricetus* (Golden hamster), *Myotis lucifugus* (Little Brown bat) and *Hipposideros armiger* (Great Roundleaf bat) in DF1 cells. Results demonstrate permissive replication of SC2 in cat, Golden hamster, and goat species, but not pig or horse, which correlated with the results of reported challenge studies. The development of this cell culture model allows for more efficient testing of the potential susceptibility of many different animal species for SC2 and emerging variant viruses.

**IMPORTANCE:** SARS-CoV-2 (SC2) is believed to have originated in animal species and jumped into humans where it has produced the greatest viral pandemic of our time. Identification of animal species susceptible to SC2 infection would provide information on potential zoonotic reservoirs, and transmission potential at the human-animal interface. Our work provides a model system to test the ability of the virus to replicate in an otherwise non-permissive cell line by transgenic insertion of the ACE2 and TMPRSS2 genes from human and other animal species. The results from our in vitro model positively correlate with animal infection studies enhancing the predicative capability of the model. Importantly, we demonstrate that both proteins are required for successful virus replication. These findings establish a framework to test other animal species for susceptibility to infection that may be critical zoonotic reservoirs for transmission, as well as to test variant viruses that arise over time.

## INTRODUCTION

The current COVID-19 pandemic is caused by the severe acute respiratory syndrome coronavirus 2 [SARS-CoV-2 (SC2)] which was first reported in Wuhan, China in late 2019. This virus most probably has its ecological reservoir in bats, and transmission of the virus to humans has likely occurred through an intermediate animal host which has not yet been identified (1, 2). Coronaviruses (CoVs) are a large family of viruses, several of which cause respiratory diseases in humans, from the common cold to more rare and serious diseases such as the Severe Acute Respiratory Syndrome (SARS) and the Middle East Respiratory Syndrome (MERS), both of which have high case fatality rates and were detected for the first time in 2002 and 2012, respectively.

CoVs are enveloped, single-stranded, positive-sense RNA viruses that belong to the subfamily *Orthocoronavirinae* within the family *Coronaviridae*, Order *Nidovirales*. The viruses are divided into four genera: alpha-, beta-, gamma- and delta-CoV based on phylogenetic and genomic structure (3, 4). All CoVs currently known to cause disease in humans belong to the alpha- or beta-CoV groups (5, 6). In addition, alpha-CoV, beta-CoV and gamma-CoV induce significant disease on various domestic animal species, including porcine transmissible gastroenteritis virus, porcine enteric diarrhea virus (PEDV), swine acute diarrhea syndrome coronavirus (SADS-CoV), and infectious bronchitis virus (IBV) in poultry (5–9). Based on sequence analysis, human coronaviruses have animal origins. The SARS-CoV, MERS-CoV, HCoV-NL63 and HCoV-229E are thought to have originated in bats, whereas HCoV-OC43 and HKU1 appear to have come from rodents (10). The 2002 SARS-CoV-1 recombined in civet cats and humans whereas the 2012 MERS-CoV appeared to have spread from bats to dromedary camels and then to humans (11–13).

The main surface protein of CoVs is the spike (S) protein that facilitates receptor binding and fusion of the viral lipid envelope with the host cell membrane. Receptor binding is facilitated by the S1 subunit while the S2 subunit is involved with fusion of the viral membrane with the cell membrane (14, 15). For these two events to occur, the S protein needs to be post-transitionally modified by two different host proteases to become activated. For SC2, furin-like proteases cleave the S protein at the S1/S2 site that contains a multiple basic amino acid motif (RRAR) that is different from SARS-CoV (16). The S protein undergoes additional cleavage at the S2’ site by the cellular type II transmembrane serine protease, TMPRSS2 (17–19). However, other proteases have been described to activate CoVs including cathepsin L, TMPRSS11A and TMPRSS11D (20–23).

SARS-CoV and SC2 utilize the angiotensin-converting enzyme 2 (ACE2) as the receptor for attachment on host cells with the S protein (14). ACE-2 is a single-pass type I transmembrane protein, with its enzymatically active domain exposed on the surface of cells in lungs and other tissues. ACE2 catalyzes the conversion of angiotensin I into angiotensin 1-9 and angiotensin II into angiotensin1-7, which are involved with vasodilation effects in the cardiovascular system (24, 25). Due to conservations of the ACE2 gene among animal species, the potential host range of SC2 is thought to be extensive.

The ACE2 and TMPRSS2 genes have homologues in many animal species (1, 22). Several species, including house cats, ferrets, and golden hamsters, have been shown to be naturally and/or experimentally infected with SC2 (26). These three species have >80% sequence similarity in their ACE2 and TMPRSS2 genes when compared to the human genes. The chicken, which does not appear to be a susceptible host, has an ACE2 homology of less than 70% to the human gene (27). However other species like pigs have a sequence similarity of >80%, but are poorly susceptible to infection. Based on previous work with SARS-CoV, the binding of S1 to ACE2 can be defined by the interaction of relatively few amino acids, and predictions of host susceptibility based on these interactions have been made (1, 28). Despite the clear importance of the binding of the spike protein to ACE2, the prediction of host susceptibility does involve other factors including the level and tissue distribution of ACE2 expression and the requirement for protease activation.

Because chickens are not susceptible to SC2 virus, and their ACE2 and TMPRSS2 protease are distinctly different from the human equivalents, we developed an avian cell line to screen the potential host range of infection of the virus through the expression the ACE2 and TMPRSS2 genes from human and animal species to provide novel insights into the receptor usage, replication and potential host range of SC2 These studies were designed to determine if the host restriction is strictly from the difference in the receptor and/or protease. One long-term goal of this work is to develop a predictive framework for improved epidemic surveillance to include protection of agriculturally relevant species and animal species that are hard to test experimentally.

## MATERIALS AND METHODS

### Viruses

The USA-WA1/2020 (BEI NR-58221, original material was provided by the US Centers for Disease Control and Prevention) isolate of SARS-CoV-2 (SC2) was obtained from BEI Research Resources Repository, National Institute of Allergy and Infectious Diseases, National Institutes of Health (29). The virus was propagated and titrated in ATCC-CCL-81 Vero cells and was utilized at 6 or 7 total passages in Vero cells. Experiments with SC2 were performed in a biosafety level-3 enhanced facility with procedures approved by the U.S. National Poultry Research Center Institutional Biosafety Committee.

### Cell lines

DF1 (avian fibroblast), Madin-Darby Canine Kidney (MDCK) and Vero (African Green monkey kidney, CCL-81) cells were seeded and propagated with standard procedures for adherent cells in flasks containing Dulbecco’s Modified Eagle Medium (DMEM) (ThermoFisher Scientific, Waltham, MA) with 10% Fetal Bovine Serum (Sigma Chemical Company, St. Louis, MO) and 1% Antimicrobial-Antimycotic (GeminiBio, Sacramento, CA). At each passage adherent cells were disassociated with trypsin (GIBCO) when at 95-100% confluence and passaged. Cells were incubated (ThermoFisher Scientific) at 37°C with 5% CO_2_. Vero cells were obtained from the International Reagent Resource (FR-243). MDCK cells were obtained from ATCC and were included because this sub-lineage was not able to support SC2 replication, therefore could serve as an additional cell line to evaluate results (14).

### Construction of transgenic cell lines using lentivirus vectors expressing human ACE2 and TMPRSS2

DF1 and MDCK cells were seeded at a density of 0.5 × 10^5^ in 500µl DMEM containing 10% Fetal Bovine Serum and 1% Antimicrobial-Antimycotic (Sigma), in one well each of a 12 well plate, and left overnight as above. Once cells reached 50-75% confluence, the media was removed and lentivirus particles were added, according to the manufacturer’s recommendations. The lentivirus contained the human ACE2 gene under control of the CMV promoter along with green fluorescent protein (GFP) also under control of a separate CMV promoter (Origene Technologies, Rockville, MD). A MOI of 20 was used for lentivirus transduction. For TMPRSS2 transduction, lentivirus particles containing the human TMPRSS2 gene under control of the CMV promoter and red fluorescent protein (RFP) gene under control of a separate CMV promoter (Gentarget, San Diego, CA), were added to achieve a MOI of 20. Polybrene (8µg/ml) was added to each transduction reaction, supplied from the manufacturers, to aid with membrane charge. Cells were incubated at 39°C for 72 hours after which media was removed and replaced with fresh media containing 10% FBS. Transduction was confirmed using an EVOS 5000 (Invitrogen, Carlsbad, CA), equipped with GFP, RFP, DAPI and transmitted light cubes, to visualize cells expressing GFP or RFP, or both. Production of DF1 or MDCK cells expressing only human ACE2 (defined as +-) or only human TMPRSS2 (defined -+), or both (defined as ++), was confirmed by RT-PCR and purification by FACS cell sorting for either green or red fluorescence. For construction of cells expressing both, the human ACE2 was first inserted and purified for GFP (99% GFP-positive) followed by human TMPRSS2 insertion and cell sorting for both RFP- and GFP-positive cells (See Supplemental figure 1). Confirmation of human ACE2 and human TMPRSS2 expression was performed by RT-PCR and western blot.

### Construction of transgenic DF1 cell lines expressing different animal ACE2 and TMPRSS2 genes using the PiggyBac transposon vector

GenBank accession numbers used to construct all species plasmids can be found in Supplemental Table 1. The ACE2 and TMPRSS2 genes from cat (*Felis catus*), horse (*Equus ferus*), domestic pig (*Sus domesticus*), goat (*Capra aegagrus*), Golden hamster (*Mesocricetus auratus*), Little Brown bat (*Myotis lucifugus*) and Great Roundleaf bat (*Hipposideros armiger*) were *de novo* synthesized into the PiggyBac® transposon expression plasmids under control of the CMV promoter (VectorBuilder Inc., Chicago, IL). As with the human genes, GFP was included for ACE2 detection and purification, and RFP was included for TMPRSS2 detection and purification. Frozen *E. coli* plasmid glycerol stocks, containing either ACE2 or TMPRSS2, were streaked onto LB agar plates (Invitrogen) containing 100 µg/mL of Carbenicillin (Sigma). Plates were incubated overnight at 34°C in an incubator/shaker (Amerex Instruments, Concord, CA). Single colonies were selected and incubated in 50 mL LB Broth, containing 100 µg/mL of Carbenicillin, with gentle agitation overnight in an incubator/shaker at 34°C (Amerex Instruments).

### Plasmid DNA Isolation

*E. coli* plasmid colonies, from overnight LB broth culture, were pelleted, by centrifugation at 4,000 x g for 10 minutes at room temperature in a tabletop centrifuge (Beckman Coulter, Pasadena, CA). Plasmid DNA was isolated from each cell pellet using the Purelink®HiPure Plasmid Maxiprep DNA Purification Kit (Invitrogen) according to manufacturer’s instructions. Purified DNA was eluted in 50 µl TE buffer. DNA was quantified using the DeNovix DS-11FX spectrophotometer/fluorometer with a Qubit™ dsDNA HS Assay Kit (Invitrogen), and stored at −20°C.

### PiggyBac Transfection with animal ACE2 or TMPRSS2

DF1 cells were seeded, at a density of 0.5 × 10^5^ in 500 µl DMEM, containing 10% Fetal Bovine Serum (FBS) and 1% Antimicrobial-Antimycotic, in one well of a 12 well plate. Cells were incubated overnight at 39°C to reach 75-90% confluence. Once cells reached desired confluence, the media was removed, and cells were washed twice with DMEM. Cells were transfected using Lipofectamine 3000 (Invitrogen) according to the manufacturer’s protocol. Transposase and Transposon DNA were added at 1:1 ratio in 10% FBS. Cells were incubated for 72 hours at 39°C, after which expression was confirmed using an EVOS 5000 as above.

### Fluorescent-activation cell sorting (FACS)

Transgenic cells expressing ACE2, TMPRSS2 or both, were grown to 90% confluence in T125 flasks. Adherent cells were trypsinized and pelleted by centrifugation at 1500 x g for 10 minutes at room temperature. Cell pellet was resuspended in phenol red free DMEM (GIBCO) containing 20% FBS, and 1% Antimicrobial-Antimycotic. The cell suspension was then strained through a 50µm cell strainer (Fisher Scientific). Cells were sorted for GFP or RFP, or both, at the University of Georgia (Athens, Georgia), Flow Cytometry Core Center, using a Beckman Coulter Moflo Astrios EQ (Beckman Coulter).

### RNA extraction and RT-PCR for human ACE2 and TMPRSS2

Total RNA was extracted from 2.5 x 10^5^ cells in one well of a 6 well plate from Vero, DF1, DF1 +-, DF1 -+, DF1 ++, MDCK, MDCK +-, MDCK -+ and MDCK ++. Once cells were 75% confluent, media was removed and 500 µl of Trizol Reagent (Invitrogen) was added to the wells then placed into 1.5 mL microcentrifuge tubes. Tubes were centrifuged at 10,000 x g for ten minutes at 4°C to remove any solids. One hundred µl of chloroform (Sigma) was added to supernatant, mixed by rapid inversion for 30 seconds, allowed to sit for 3 minutes, and centrifuged at 10,000 x g for 15 minutes at 4°C. The aqueous phase was then removed and added to an equal amount of 100% Ethanol (Sigma). Final RNA extraction was carried out using the ZYMO Direct-zol Mini-Prep Plus Kit (Zymo Research, Irvine, CA) per manufactures instructions.

Superscript 4 Reverse Transcriptase (Invitrogen) was used according to manufacturer’s instructions. One µl of 2 µm gene specific primer and 11 µl of RNA were used for all reactions. Gene specific first strand primers used were: human ACE2 5’ GGA TCC TAA AAG GAG GTC TGA ACA TCA TCA 3’ and human TMPRSS2 5’ GAA TCG ACG TTC CCC TGC AG 3’. Two µl of cDNA template was used for all cell lines. Reactions were conducted using NEB Phusion Hi Fi Polymerase (New England Biolabs, Ipswich, MA). Reactions were comprised of 4 µl 5X Phusion Buffer, 0.4 µl 10 mM DNTPs, 1 µl of Forward and Reverse Primer, 2 µl of cDNA, 0.6 µl of DMSO, 0.2µl of DNA polymerase, and 11 µl of ultrapure water (Invitrogen). Primers used for human ACE2 PCR were Forward 5’ CTA GCT GTC AAG CTCTTC CTG GCT C 3’ and Reverse 5’ GGA TCC TAA AAG GAG GTC TGA ACA TCA TCA 3’. Reaction conditions were 98°C for thirty seconds, followed by 35 cycles of 98° for ten seconds, 68°C for thirty seconds and 72°C for one minute, after which a final extension of ten minutes at 72° was added.

Primers for human TMPRSS2 were Forward 5’ GGA AAA CCC CTA TCC CGC AC3’ and Reverse 5’ GAA TCG ACG TTC CCC TGC AG 3’. Annealing temperature for reactions was 66°C and all other conditions were identical to human ACE2. PCR products were visualized on 1% agarose gel (Bio-Rad Laboratories, Hercules, CA) containing SYBR Safe (Invitrogen) using a documentation system (Syngene International Ltd, Bengaluru, India).

### RNA extraction and RT-PCR for animal species ACE2 and TMPRSS2

Total RNA was extracted as above. Superscript 4 Reverse Transcriptase (Invitrogen) was used according to manufacturer’s instructions. One µl of 2 µm gene specific primer and 11 µl of RNA were used for all reactions. Gene specific first strand primers used were: universal (except chicken) ACE2 5’ TCC AAG AGC TGA TTT TAG GCT TAT CC 3’ and universal (except bat and chicken) TMPRSS2 5’ CTG TTT GCC CTC ATT TGT CGA TA3’. Bat TMPRSS2 first strand primers were: 5’ CAA AGT GAC CAG AGG ACC G 3’. Chicken ACE2 first strand primer 5’AGC CAA TGG ATC TGC CAG AA 3’ and chicken TMPRSS2 first strand primers 5’ TCT GCC AGG CCA CAA GTA GG 3’. Two µl of cDNA template was used for all cell lines. Reactions were conducted using NEB Phusion Hi Fi Polymerase (New England Biolabs, Ipswich, MA). Reactions were comprised of 4 µl 5X Phusion Buffer, 0.4 µl 10 mM DNTPs, 1 µl of Forward and Reverse Primer, 2 µl of cDNA, 0.6 µl of DMSO, 0.2µl of DNA polymerase, and 11 µl of ultrapure water (Invitrogen). Primers used for animal (except chicken) ACE2 PCR were Forward 5’ CTC TTT CTG GCT CCT TCT CAG CTT 3’ and Reverse 5’ TCC AAG AGC TGA TTT TAG GCT TAT CC 3’. Chicken ACE2 primers were Forward 5’ACG CTA GCC GCT TCT CAC TAG C 3’ and Reverse 5’AGC CAA TGG ATC TGC CAG AA 3’. Reaction conditions were 98°C for thirty seconds, followed by 35 cycles of 98° for ten seconds, 68°C for thirty seconds and 72°C for one minute, after which a final extension of ten minutes at 72° was added.

Universal primers for animal TMPRSS2 (except bat and chicken) were Forward 5’ ATG GCT TTG AAC TCA GGG TC 3’ and Reverse 5’ CTG TTT GCC CTC ATT TGT CGA TA 3’. Bat TMPRSS2 primers were Forward 5’ CAG GGA TTT TGA GAC AAT CTT TCA T 3’ and Reverse 5’ CAA AGT GAC CAG AGG ACC G 3’. Chicken specific TMPRSS2 primers were Forward 5’TGT TAC CAG AGG ACC TCC GC 3’ and Reverse 5’ TCT GCC AGG CCA CAA GTA GG 3’. Annealing temperature for reactions was 66°C and all other conditions were identical to animal ACE2. PCR products were visualized on 1% agarose gel (Bio-Rad Laboratories, Hercules, CA) containing SYBR Safe (Invitrogen) using a documentation system (Syngene International Ltd, Bengaluru, India). All primers used in these studies are listed in Supplemental Table 2.

### Detection of human ACE2 and TMPRSS2 protein expression by western blot, and immunohistochemistry to detect SC2

Total cellular protein was extracted from cells seeded into one well of a six well plate in 10% FBS as above. Once cells reached 75% confluence media was removed and cells were washed twice with 1X PBS. One hundred µl of 2X Laemmli buffer, containing 2-mercaptoethanol, was added to the cells and collected into 1.5 ml microcentrifuge tubes. The cells were then boiled for 7 minutes and vortexed. Fifteen µg of each protein sample and 5 µl of Page Ruler Plus (Invitrogen) was loaded onto a Bio Rad Mini-Protean Precast TGX gel and separated for one hour at 100 Volts. The separated proteins were transferred to a 0.2 µM nitrocellulose membrane (Bio Rad) at 100V for 1 hour as previously described (30). Unbound proteins binding sites were blocked with 3% non-fat milk in 1X PBS for 1 hour at room temperature with gentle rocking. The blot was washed 3 times, for five minutes, with 1 X Tris Buffered Saline (TBS), pH 7.4, containing 0.05% tween-20 (TBST). The blot was then incubated overnight at 4°C in primary antibody diluted 1:1500 in TBS. Primary monoclonal antibodies included mouse anti-human ACE2 (Origen), rabbit anti-human TMPRSS2 (Abcam, Cambridge, UK) and mouse anti-beta actin (Invitrogen). The blot was washed as before, incubated for 1 hour, at room temperature, in secondary antibody diluted 1:20,000 in TBS with gentle rocking. Secondary antibodies included rat anti-mouse IgG1 HRP (Southern Biotech, Birmingham, AL), and mouse anti-rabbit IgG1 HRP (Southern Biotech). After incubation, the blot was washed 3 times as above in TBST. Pierce ECL substrate (Fisher) was added to the blot for 1 minute and excess was removed by gentle wicking. The blot was placed into an x-ray cassette and exposed to x-ray film (Fisher) for 1 minute, developed and fixed (Kodak).

For immunohistochemistry of SC2 replication, cells were seeded into an I-Bidi 8-well chambered slide (Fisher) at a density of 4 × 10^4^ in 500 ul DMEM containing 10% FBS and grown overnight as above. When cells reached 75% confluence the media was removed, and virus was added at MOI of 1 as above. After 48 hours, the media was removed and cells were fixed for 5 minutes at 4C in 1:1 ice cold ethanol:methanol. Cells were then washed twice with cold PBS as above. Cells were blocked as above for one hour at room temperature then washed 3 time with TBS. Primary antibodies against SC2 included rabbit anti-Nucleoprotein MAb (Origene) and rabbit anti-Spike MAb (Origene), diluted as above, were added for 1 hour at room temperature. Cells were washed 3 times with PBS and incubated in the secondary antibody, goat anti-rabbit IgG H&L (Alexa Fluor^®^ 555) (ABCAM) diluted 1:20,000 in TBS, for one hour at room temperature. Cells were then washed 3 times with PBS and counterstained with DAPI (Invitrogen) for 5 minutes. Cells were washed 3 times with PBS then allowed to air dry. Once dry, cells were mounted with ProLong™ Gold Antifade Mountant (Fisher) and sealed with glass coverslips after 24 hours. Immunofluorescence was visualized with an EVOS 5000 (Invitrogen).

### Comparison of SARS-CoV-2 replication dynamics among cell lines

Cell lines were tested for virus replication by inoculating them with SC2 at an MOI of 1 added directly when cells were approximately 70-90% confluent in 6 well plates. For each cell line, media was removed from three wells and 0.4 ml of virus was added. The same volume of sterile medium was added to wells on each plate to serve as a sham inoculated control. The plates were incubated for 1 hr at 37°C, 5% CO_2_ to allow virus to adsorb to the cells. Each well was washed 3-times with sterile PBS prewarmed at 37°C to remove unbound virus. Finally, 3 ml growth medium was added to each well and the cells were incubated at 37°C with 5% CO_2_. Supernatant (0.2mL) was collected from each well individually at 6, 12, 24, 36, 48 and 72 hours post inoculation (hpi) for detection of replicating virus by RT-PCR, and detection of cytopathic effect. After 72 hpi, plates were frozen and thawed at −80C (3x total) and 400 ul of cell culture supernatant was transferred onto fresh cell cultures as above for a pass 2.

### Quantitative real-time RT-PCR to detect SARS-CoV-2

Quantitative RT-PCR was utilized to detect and determine virus titers in cell culture supernatants. RNA was extracted with the Ambion Magmax kit (ThermoFisher). The US Centers for Disease Control N1 primers and probe for SARS-CoV-2 were used with the AgPath ID one-step RT-PCR kit (31). The cycling conditions for the RT step were modified to accommodate the recommended kit conditions. A standard curve of RNA from titrated SARS-CoV-2 virus stock was run in duplicate to establish titer equivalents of virus.

### TMPRSS2 genetic analysis

TMPRSS2 gene sequences from animal species were obtained from GenBank. Sequences were aligned with Clustal V (Lasergene 10.0, DNAStar, Madison, WI), and protein architecture derived from The National Center for Biotechnology (www.ncbi.nih.gov).

### Statistical analysis

Viral titers at 48 hpi were compared with the two-way ANOVA with Tukey multiple comparison (Prism 9.1.0 GraphPad Software, San Diego, CA). Different lower case letters indicate statistical significance between compared groups. All statistical tests used P < 0.05 as being statistically significant.

## RESULTS

### Development of DF1 and MDCK cell lines expressing human ACE2 and TMPRSS2

These studies were designed to transgenically introduce the human receptor and protease used by SC2 into the avian non-permissive cell line, DF1, and MDCK, to test requirements for replication competence and establish a model for infection potential. A lentivirus approach was used to deliver the human ACE2 and human TMPRSS2 genes, under control of the CMV promoter. The lentivirus constructs co-expressed GFP (ACE2) and/or RFP (TMPRSS2) to allow FACS sorting for purification of cells containing each target gene or both genes (Supplemental Figure 1A, B, C). Positive DF1 and MDCK cells were demonstrated expressing either the human ACE2 gene or human TMPRSS2 gene alone, or both, based on microscopy and two-color cell sorting (Figure 1, Supplemental Figure 2). Detection of the inserted genes was confirmed with RT-PCR using primers specific for the human and chicken genes (Figure 2A and B). Expression of human ACE2 and human TMPRSS2 protein in DF1 ++ and MDCK ++ cells was confirmed via western blot (Figure 2C).

**Figure 1.**
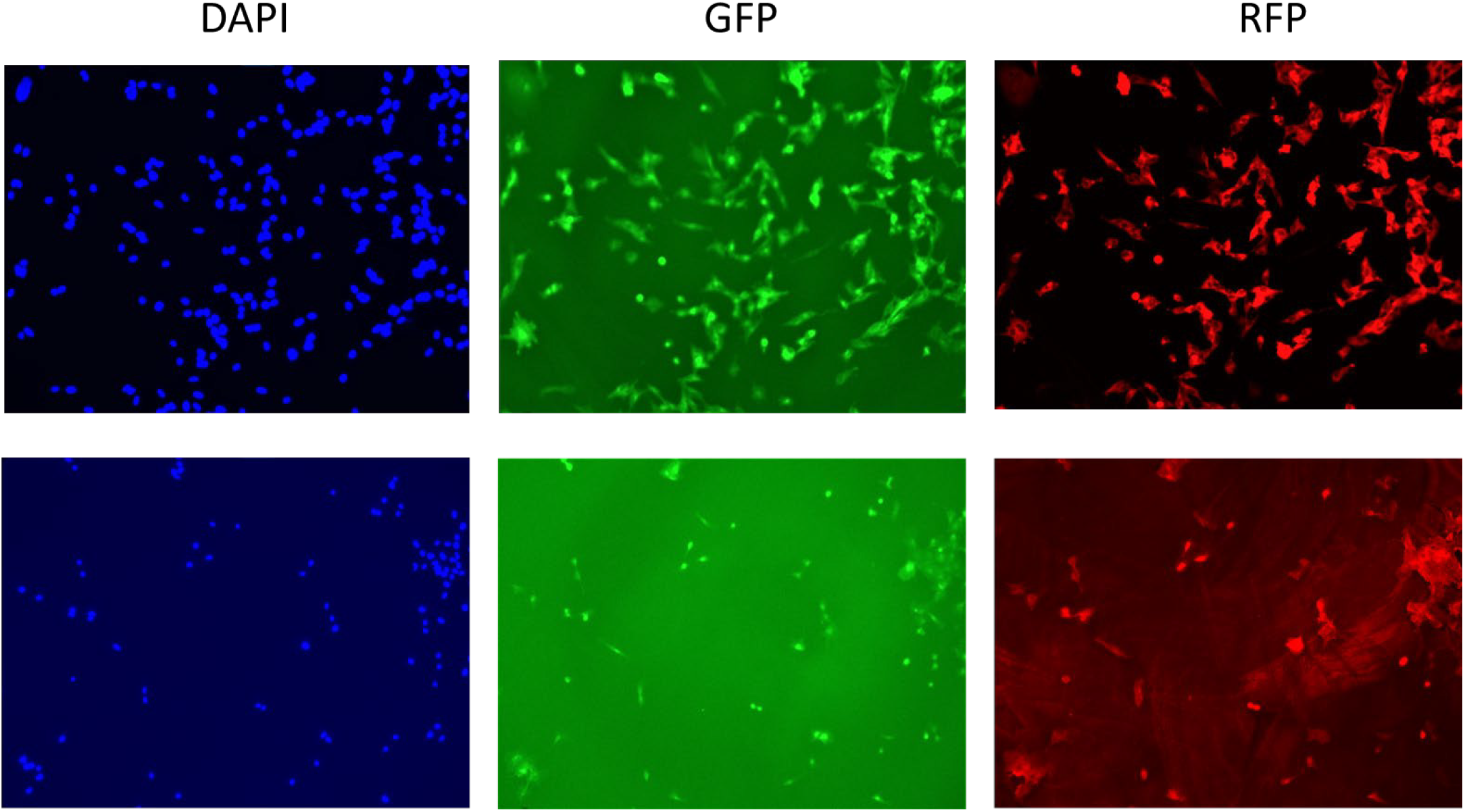
DF1 and MDCK cells expressing the human ACE2 (with GFP marker) and TMPRSS2 (with RFP marker) genes. DF1 and MDCK cells were transduced with lentivirus containing the human ACE2 gene and cells were FACS purified based on GFP expression. Lentivirus containing the humanTMPRSS2 gene was then transduced into the human-ACE2 expressing DF1 and MDCK cells. Following two-color FACS for GFP and RFP expressing cells, dual positive cells were grown for 48 hours in an 8-chamber glass slide. Fluorescence was captured on an EVOS M5000 with added DAPI nuclear stain (blue) GFP and RFP.

**Figure 2.**
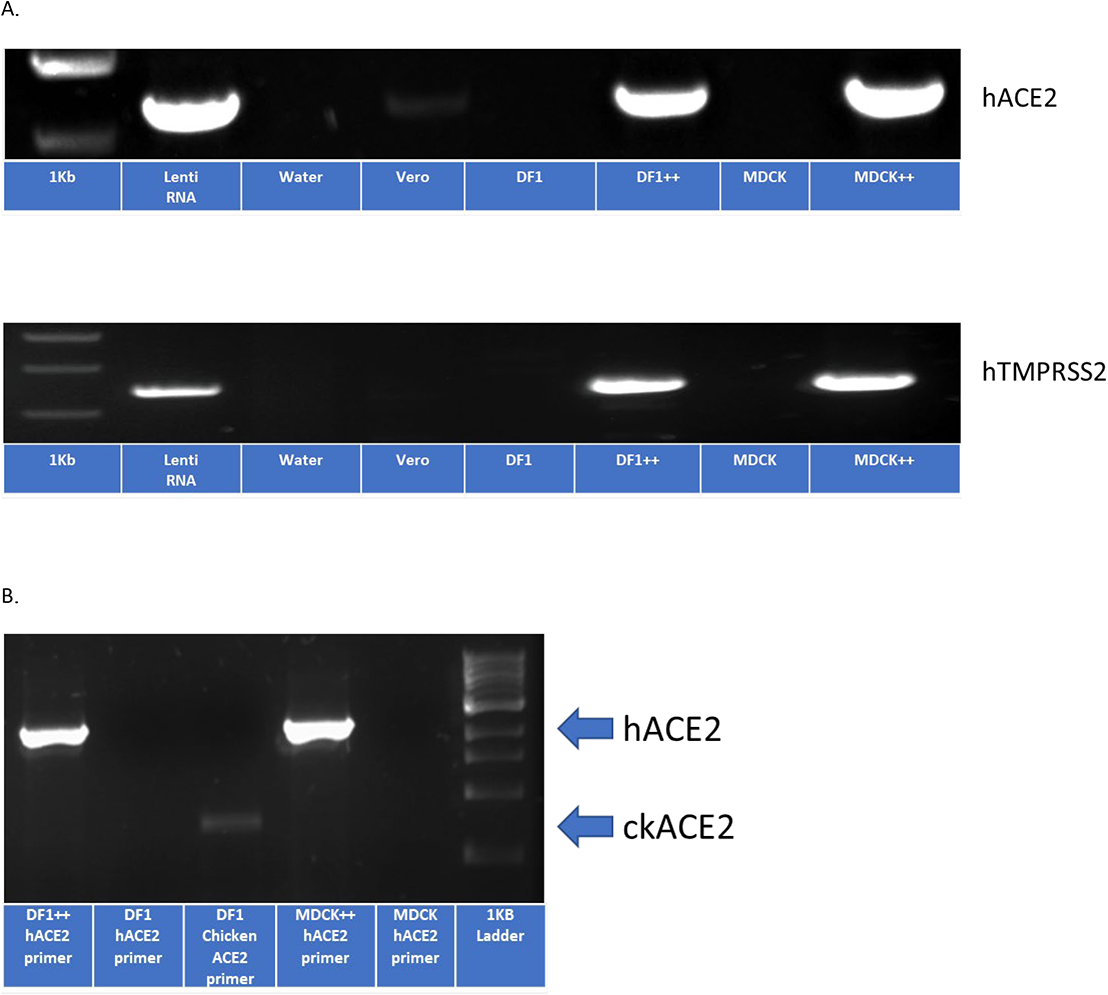
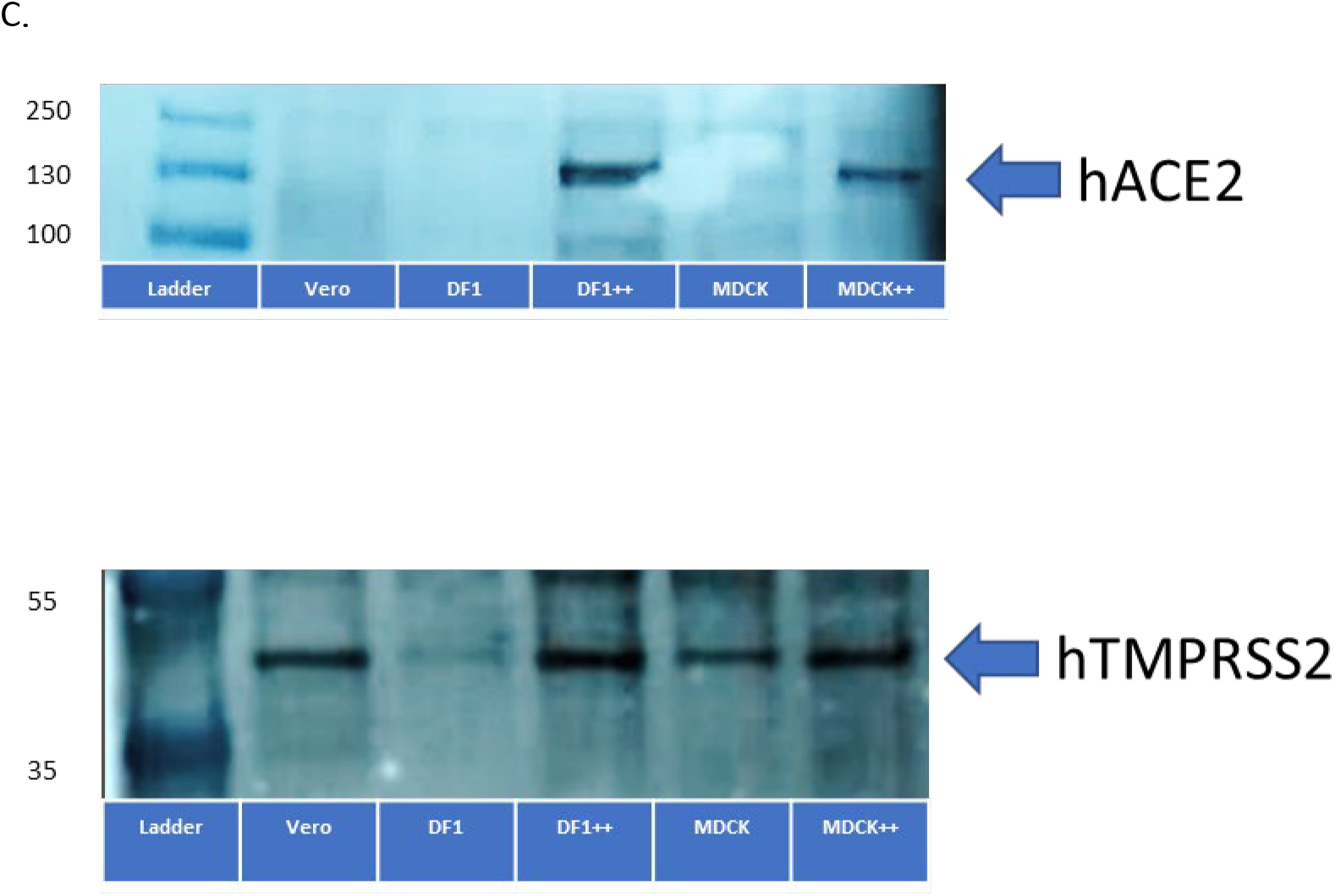
Detection of human ACE2 and human TMPRSS2 expression in DF1 ++ and MDCK ++ cells. (A) DF1, DF1 ++, MDCK, MDCK ++ and Vero cells were grown at 37C in 5% CO_2_. After 72 hours, RNA was extracted and primers specific for human ACE2 and human TMPRSS2 were used with RT-PCR to confirm expression in DF1 ++ and MDCK ++ cell lines. (B) Differential expression of human and chicken ACE2 in DF1, DF1 ++, MDCK, and MDCK ++ cell lines with primers specific for both. (C) Fifteen micrograms of protein were extracted from each cell line and separated by SDS-PAGE. Following transfer to nitrocellulose, membranes were probed by western blot using rabbit monoclonal antibodies to the human ACE2 and TMPRSS2 proteins.

### Comparison of SARS-CoV-2 replication dynamics in DF1 and MDCK cell lines expressing human ACE2 and/or TMPRSS2

Growth curves for all three cell lines (Vero, DF1, and MDCK) expressing only human ACE2 (+-), only human TMPRSS2 (-+), or both (++) are shown in Figure 3. No increase in virus titer was demonstrated in wild type DF1 or MDCK, or the DF1 and MDCK cells expressing singe gene constructs with human ACE2 or human TMPRSS2 (Figure 3A). In contrast, virus replication was observed in Vero (positive control), and the DF1++ and MDCK ++ cells. Virus growth was exponential until approximately 36 hours post infection and was statistically higher in these cells than others tested. Virus titers reached similar levels of approximately 10^5.6^ TCID_50_ in these three cell lines, and demonstrated a requirement for expression of both the receptor and the protease. We next passaged the 72 hour sample from each cell line after a freeze thaw cycle onto a subsequent plate of the same cells (Figure 3B). No evidence of increased replication was seen in cell lines that did not demonstrate signs of virus replication during the first passage. In contrast, the Vero, DF1++, and MDCK ++ passage 1 samples contained enough virus to induce infection and replication on passage 2, although the growth curves displayed a more linear increase in virus titer over time compared to passage 1 inoculated cells.

**Figure 3.**
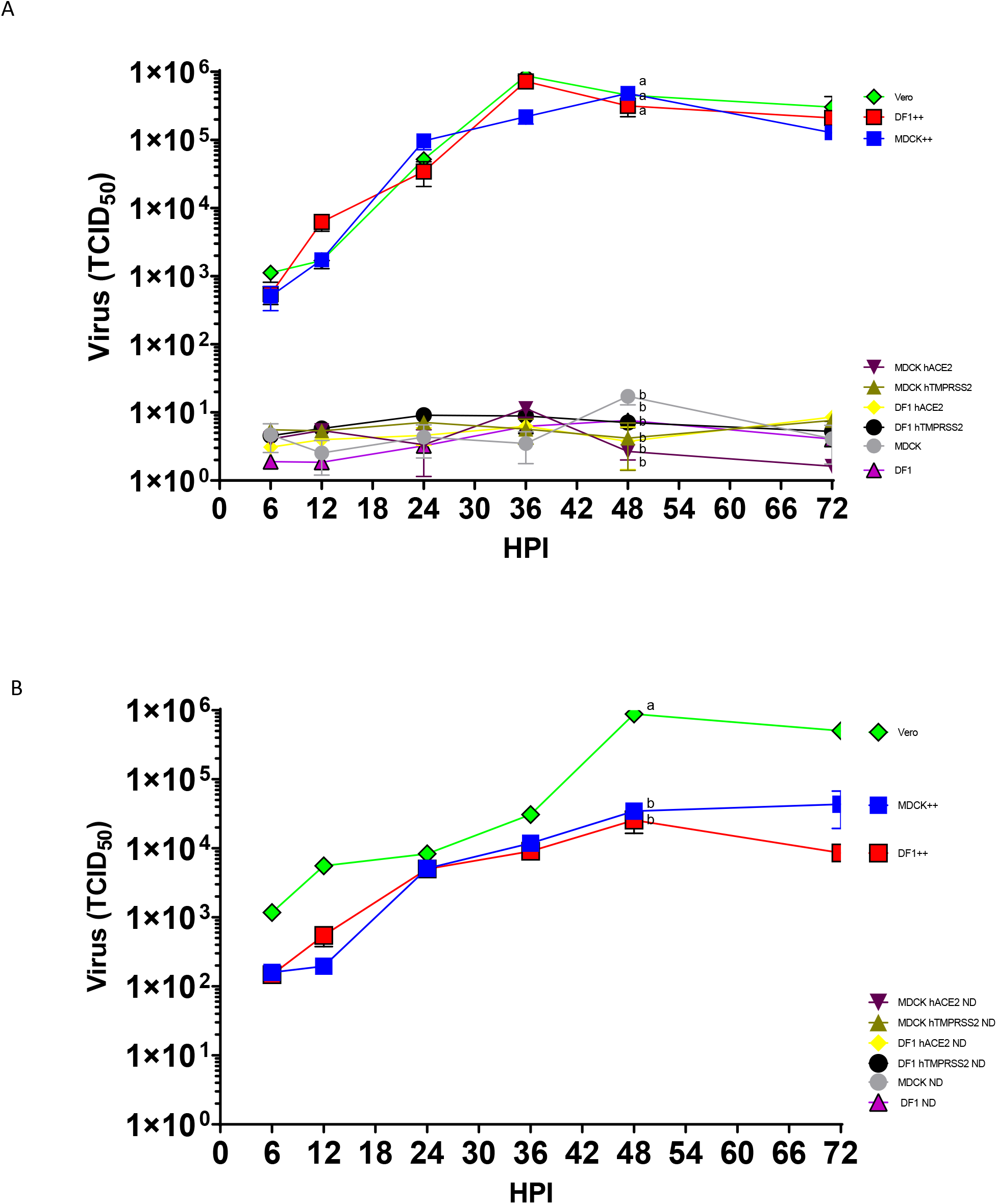
Growth of SARS-CoV-2 on DF1 and MDCK cells expressing either human ACE2, human TMPRSS2, or both (++). (A) DF1, DF1 expressing human ACE2, DF1 expressing human TMPRSS2, DF1 expressing both human ACE2 and TMPRSS2 (++), MDCK, MDCK expressing human ACE2, MDCK expressing human TMPRSS2, MDCK expressing both human ACE2 and TMPRSS2 (++), and Vero cells were inoculated with SC2 at multiplicity of infection (MOI) of 1. At time points indicated, supernatant samples were taken for RNA extraction and determination of viral titers by RT-PCR. The values shown are mean +/- standard deviation of triplicate samples. Two-way analysis of variance with Tukeys multiple comparison test was performed on titers at 48 hours post inoculation to determine the statistical difference in virus titer between the cell lines. Lines with different lowercase letters indicate differences (*p*<0.05). (B) Pass 2 of virus from cell culture lines expressing human ACE2, TMPRSS2, or both. After 72 hours of growth, supernatants of pass 1 were transferred onto fresh monolayers of cells, allowed to absorb for 1 hour and removed. Fresh media was added and samples were taken at time points indicated to determine virus titer by RT-PCR. Statistical analysis was performed at 48 hours post inoculation. ND=Not detected.

### Comparison of cytopathic effects and detection of virus in cell lines expressing human ACE2 and TMPRSS2

The appearance of CPE and confirmation of virus protein inside of the cell lines was performed via light microscopy and immunohistochemistry with antibodies against the SC2 spike and nucleoprotein. Neither CPE nor virus could be detected in cells without virus (Figure 4) or in the DF1 and MDCK inoculated cells. Likewise, cell lines containing the singular insertion of either the human ACE2 or TMPRSS2 did not exhibit CPE or positive viral staining (data not shown). Vero, DF1++, and MDCK++ demonstrated syncytia formation with loss of cell confluence. The monolayer also deteriorated by 72 hpi and CPE correlated with detection of high levels of expression of the viral spike and nucleoprotein by immunostaining at 48 hpi.

**Figure 4.**
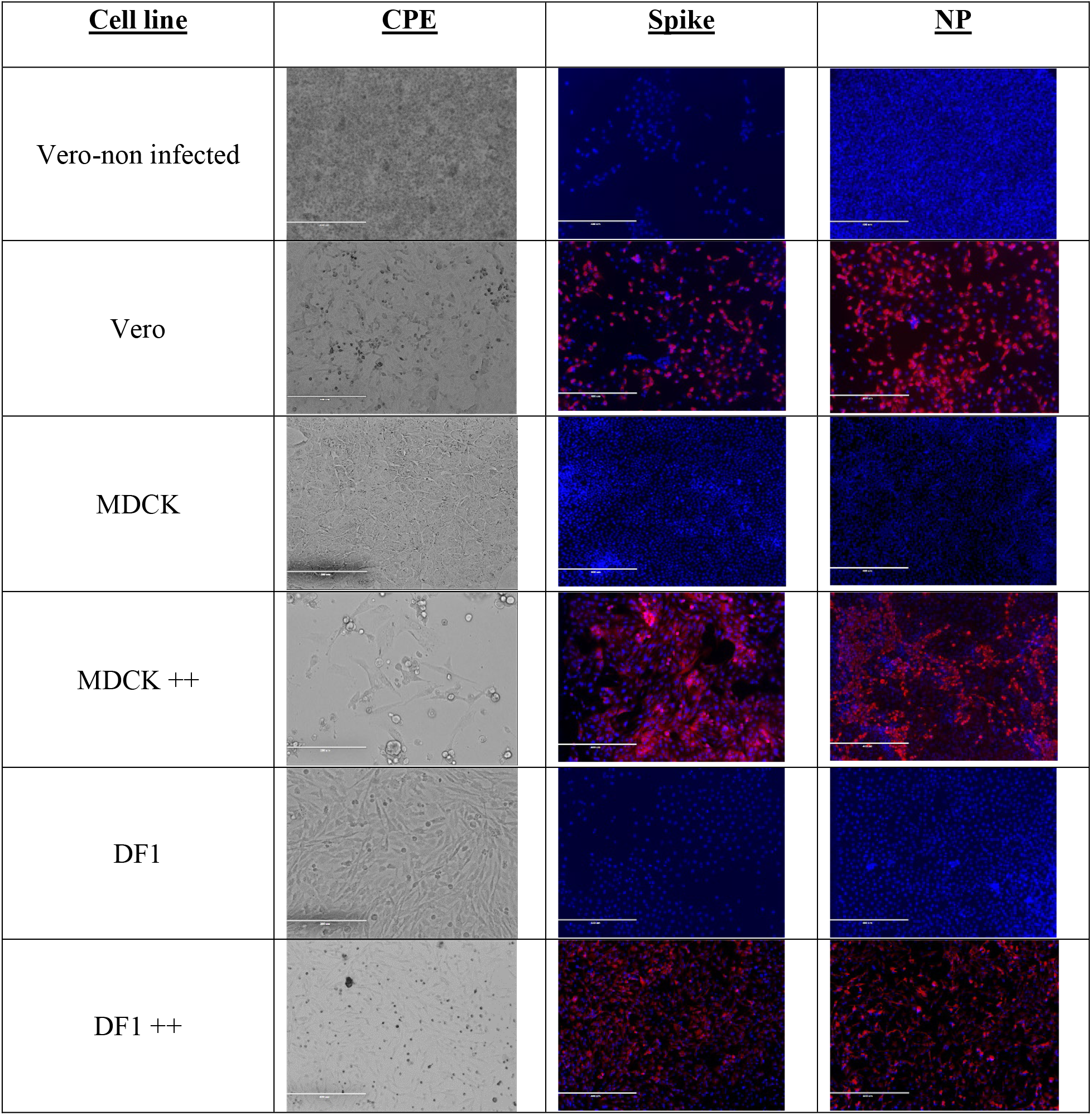
SARS-CoV-2-induced cytopathic effect and viral detection by immunohistochemistry in cells expressing human ACE2 and TMPRSS2. Vero, DF1, DF1 expressing both human ACE2 and TMPRSS2 (++), MDCK, and MDCK expressing both human ACE2 and TMPRSS2 (++) were grown at 37C in 5% CO_2_ on glass chamber slides. Cells were inoculated with SC2 at MOI of 1. At 48 hours post inoculation monolayers were examined for cytopathic effect and detection of virus with rabbit monoclonal antibodies against SC2 spike and nucleoprotein. Cells were washed 3 times with PBS and incubated in the secondary antibody, goat anti-rabbit IgG H&L (Alexa Fluor® 555) for one hour at room temperature. Cells were then washed counterstained with DAPI. Immunofluorescence was visualized with an EVOS 5000.

### Development of cell lines expressing ACE2 and TMPRSS2 from different animal species

Having demonstrated a model of virus replication in the non-permissive avian DF1 cell line with insertion of the human ACE2 and TMPRSS2 genes, we next developed cells lines expressing other species ACE2 and TMPRSS2 to screen for potential animal hosts that could support replication. The ACE2 and TMPRSS2 genes from house cat, goat, golden hamster, horse, pig, Little Brown bat, and Great Roundleaf bat were *de novo* constructed in the PiggyBac transposon system and transfected into DF1 cells. Purification of cells with green/red fluorescence was used as with the lentivirus system. As demonstrated in Figure 5, RT-PCR confirmed expression of animal ACE2 and TMPRSS2 in DF1 cells from FACS-sorted cells.

**Figure 5.**
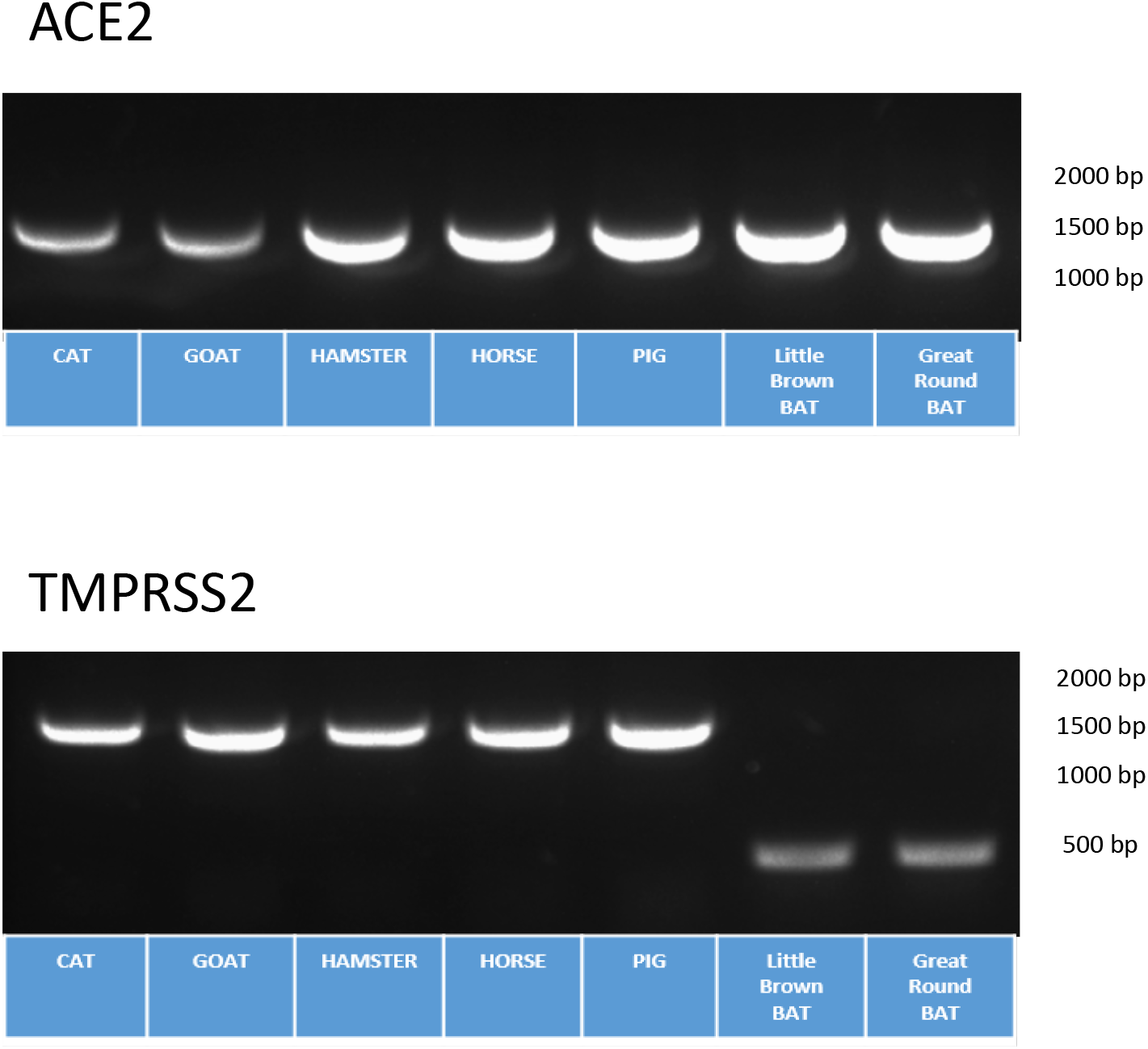
Transgenic DF1 cells expressing different animal species ACE2 and TMPRSS2 genes. (A) DF1 cells were transfected with PiggyBac® plasmid containing the ACE2 and TMPRSS2 genes from house cat (*Felis catus*), horse (*Equus ferus*), domestic pig (*Sus domesticus*), goat (*Capra aegagrus*), Golden hamster (*Mesocricetus auratus*), Little Brown bat (*Myotis lucifugus*) and Great Roundleaf bat (*Hipposideros armiger*). Cells were first created with the animal ACE2 gene and FACS purified based on GFP expression. The animal TMPRSS2 gene was then transfected into the DF1 cells expressing the animal ACE2 gene. Two-color FACS was performed based on GFP and RFP expression. Transgenic cells expressing animal ACE2 and TMPRSS2 were grown at 37C in 5% CO_2_. After 72 hours, RNA was extracted and primers specific for the animals ACE2 and animal TMPRSS2 were used with RT-PCR to confirm animal species ACE2 and TMPRSS2 expression in DF1 cells.

### SARS-CoV-2 replication in cells expressing animal ACE2 or TMPRSS2

The replication kinetics of SC2 virus in DF1 cell lines expressing the ACE2 and TMPRSS2 genes from the different animal species was determined. Results demonstrate that the SC2 virus could replicate to high levels in DF1 cell lines expressing the ACE2 and TMPRSS2 genes from cat, goat and golden hamster (Figure 6A). Virus titers reached similar levels of approximately 10^5.1^ to 10^5.8^ TCID50 at 36 hours post infection in these lines, which was similar to that observed in the Vero control cells. No virus replication was observed in the cells expressing the receptor and protease from pig or horse species. Both bat species demonstrated initial gains in virus titers, between 10^3.3^ and 10^3.9^ TCID_50_ at 12 hours post infection that did not increase after this time. The 72 hpi sample from all cell lines were passaged onto a subsequent plate of cells. Passage 2 results indicate viral infection and replication from plates containing the cat, goat and golden hamster animal cell lines (Figure 6B). As observed previously, a linear shaped curve in virus replication was observed in passage 2. Neither the pig nor the horse cell lines had evidence of virus replication in passage 2. The samples from the two bat species cell lines also had no evidence of replication on passage 2.

**Figure 6.**
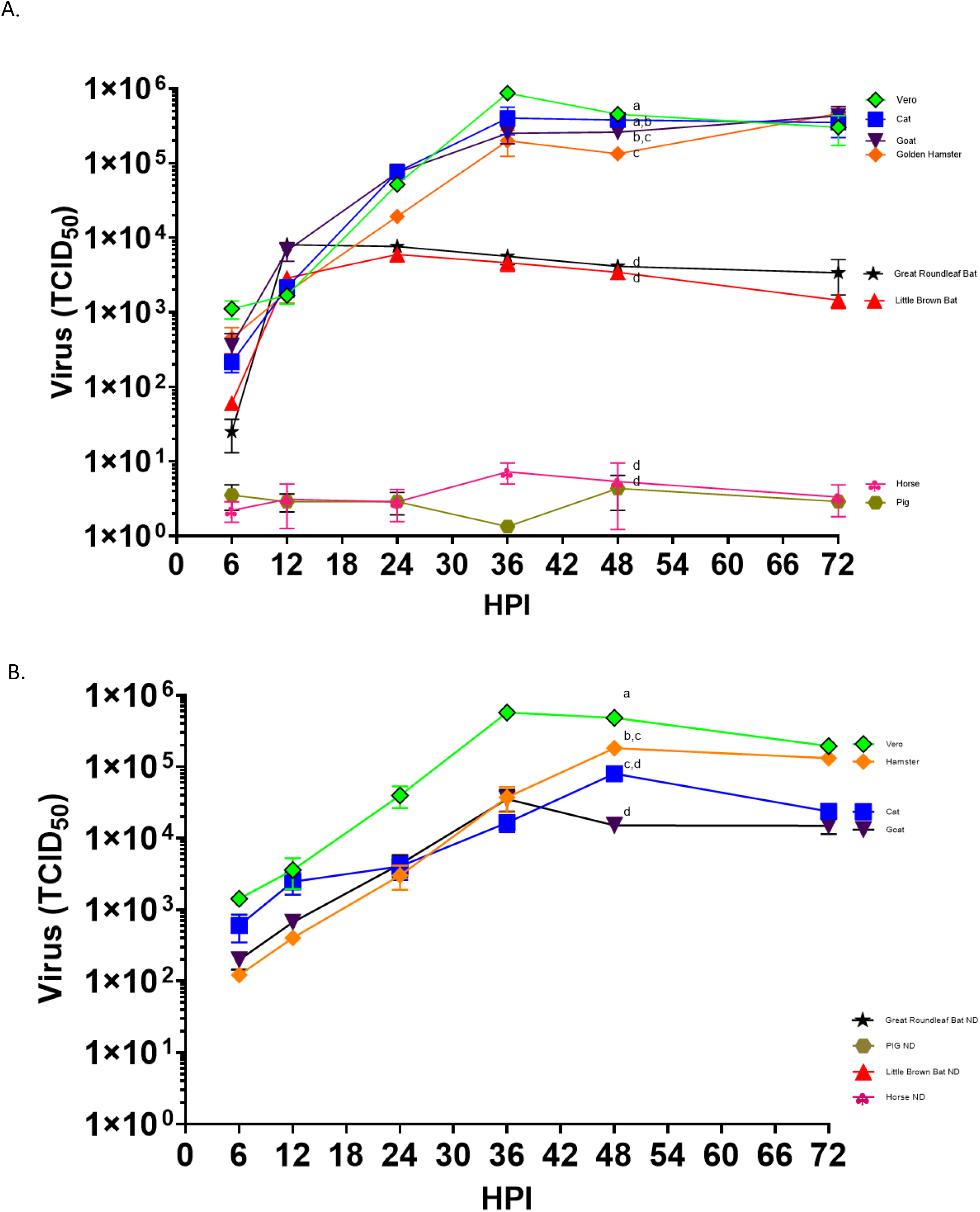
Growth of SARS-CoV-2 in DF1 cells expressing ACE2 and TMPRSS2 from different animal species. (A) DF1 cells expressing cat, horse, pig, goat, Golden hamster, Little Brown bat, and Great Roundleaf bat were inoculated with SC2 at multiplicity of infection (MOI) of 1. At time points indicated, supernatant samples were taken for RNA extraction and determination of viral titers with RT-PCR. The values shown are mean +/- standard deviation of triplicate samples. Two-way analysis of variance with Tukeys multiple comparison test was performed on titers at 48 hours post inoculation to determine the statistical difference in virus titer between the cell lines. Lines with different lowercase letters indicate differences (p<0.05). (B) Pass 2 of virus from cell culture lines animal species ACE2 and TMPRSS2. After 72 hours of growth, supernatants of pass 1 were transferred onto fresh monolayers of cells, allowed to absorb for 1 hour and removed. Fresh media was added and samples were taken at time points indicated to determine virus titer with RT-PCR. Statistical analysis was performed at 48 hours post inoculation. ND=Not detected.

Sequence analysis of available TMPRSS2 sequence data for human and animal species demonstrated a truncation at the 5’ end of the bat protein compared to human or other animals (Supplemental Figure 3). The human protein has 492 amino acids (AA), whereas the Little Brown bat contains 243 AA and Great Roundleaf bat has 384 AA. It is not clear if the bat sequences available in GenBank were incorrectly annotated and are not representative of the complete protein, and that the bat species TMPRSS2 tested here may not be functional due to the missing the N-terminal portion of the protein. The Little Brown bat open reading frame begins at human amino acid position 255, and the Great Roundleaf bat begins at human position 113. Interestingly, Brandts bat (*Myotis brandtii*) contained a protease similar to human and other animals.

### Comparison of cytopathic effects and detection of virus in cell lines expressing animal ACE2 and TMPRSS2

As before, detection of virus was observed via CPE and immunostaining of transgenic cell lines. As demonstrated in Figure 7, we detected cytopathic effects in cell lines that supported growth of the virus, including the ones expressing the cat, goat and golden hamster genes. We also observed CPE in both the cell lines expressing the bat genes which appeared more rapidly in the Great Roundleaf bat cell line compared to the Little Brown bat cell line. Staining for viral proteins was greatest in cells expressing cat, goat or golden hamster transgenes. Interestingly, we did observe positive staining in the bat species cells, however, it was visibly reduced compared to the other positive cell lines. We did not observe either CPE or viral staining in the cell lines expressing pig and horse genes.

**Figure 7.**
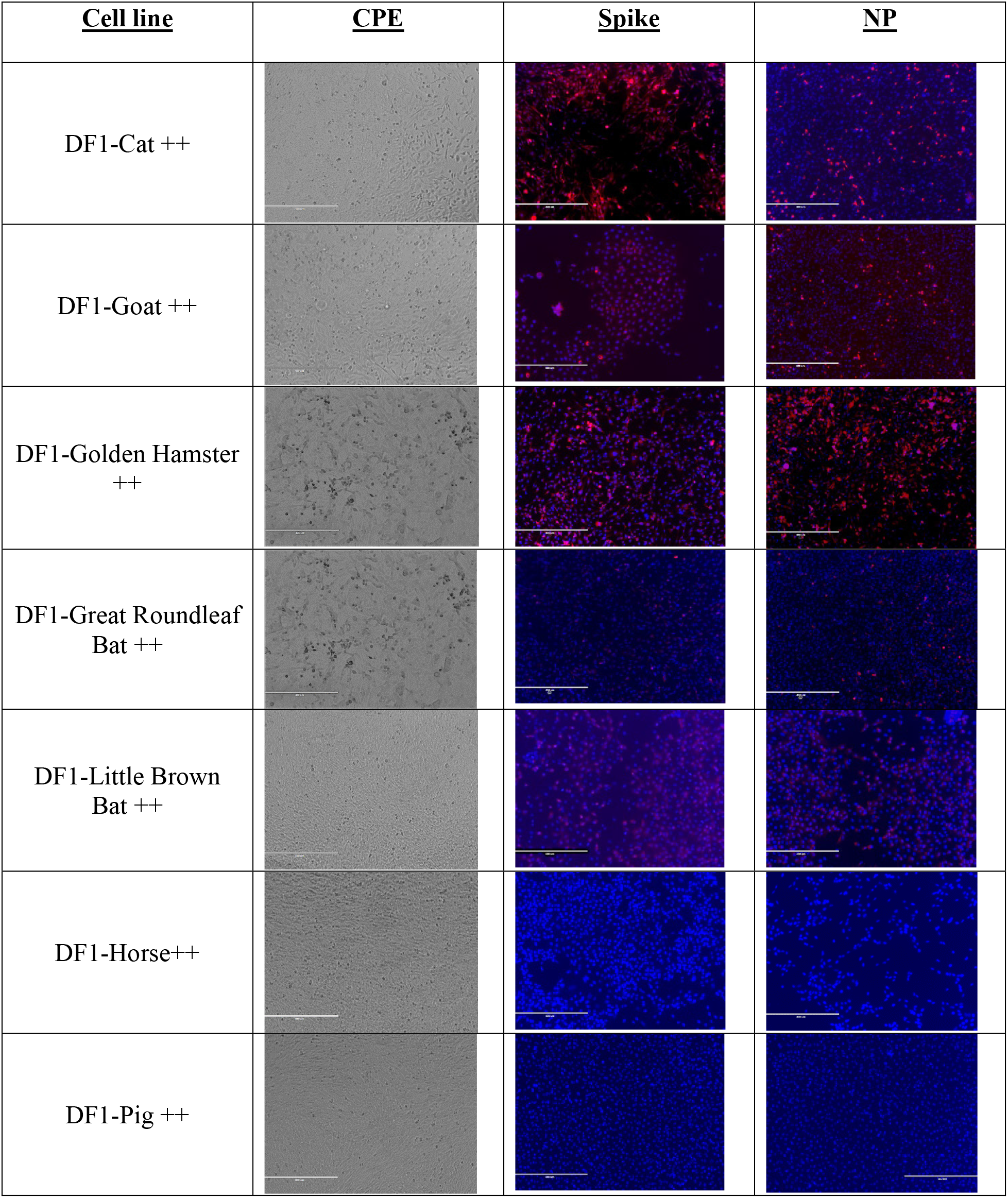
SARS-CoV-2 induced cytopathic effect and viral detection by immunohistochemistry in DF1 cells expressing animal species ACE2 and TMPRSS2. DF1 cells expressing animal ACE2 and TMPRSS2 were grown at 37C in 5% CO_2_ on glass chamber slides. Cells were inoculated with SC2 at MOI of 1. At 48 hours post inoculation monolayers were examined for cytopathic effect and detection of virus with rabbit monoclonal antibodies against SC2 spike and nucleoprotein. Cells were washed 3 times with PBS and incubated in the secondary antibody, goat anti-rabbit IgG H&L (Alexa Fluor® 555) for one hour at room temperature. Cells were then washed counterstained with DAPI. Immunofluorescence was visualized with an EVOS 5000.

## DISCUSSION

Several cell lines and organoids are currently in use or have been developed to study SC2 replication. Besides Vero cells, Caco-2, Calu-2, and Hek293T cells, human lung, kidney, liver and blood vessel organoids have been demonstrated to be permissive for virus growth (37–45). However, because these systems can naturally be infected, they are not useful for testing host susceptibility to the virus. Previous research done in our laboratory and by others clearly demonstrate that poultry and other bird species cannot support replication of the virus (27, 46, 47). We hypothesized that avian cell lines could become permissible to infection if they expressed a suitable ACE2 receptor and produced high enough levels of a protease that could activate SC-2. It is worth mentioning that proteolytic cleavage of the S protein at the S1/S2 interface was assumed to be provided by furin-like enzymes naturally present in the DF1 or MDCK cell lines.

The SC2 utilizes the ACE2 protein as the primary receptor for entry into host cells and the TMPRSS2 protease has been shown to be critical for cleavage/activation of the spike protein (32, 33). In these studies, the transgenic insertion of the human ACE2 and TMPRSS2 genes conferred virus attachment and replication ability in the non-permissive avian DF1 cell lines and MDCK cell lines. The results also demonstrated that single expression of either the human receptor or the protease was not sufficient to allow for virus replication in these cell lines, either through a lack of attachment or spike protein activation. These studies also demonstrate DF1 cells expressing the ACE2 and TMPRSS2 genes from different animal species can be used as an *in vitro* predictive model for virus replication. Wild type DF1 cells are normally incapable of supporting SC2 replication; however, expression of the receptor and protease genes from human, cat, goat and golden hamster allowed virus replication. This *in vitro* model correlates with the known natural or experimental susceptibility of three of these species and supports its use as a predictive model. The surprising result is the potential susceptibility of goats. Goats have not been known to be naturally or experimentally infected at this time, but one study has previously suggested that SC2 can infect HEK cells that are expressing goat ACE2 (53).

Multiple studies have looked at experimental inoculation in pigs, swine cell lines, and in cell lines where the swine ACE2 gene has been expressed with mixed results. Three different experimental challenge studies with swine were conducted with 2 studies showing no infection and a third showing only a small number of pigs infected after challenge (46, 47, 50–52). Two swine cell lines, swine testicular and porcine kidney cells, were also found to develop CPE after several passages of virus. Multiple studies have also used different mammalian cell lines and transfected them with the swine ACE2 gene to allow for transient expression of the gene, and most found that SC2 or a SC2 pseudovirus could attach to and express protein in the cell as measured by several different methods (2, 4, 44, 53). This predictive data based on ACE2 data from some species, such as swine, suggest susceptibility to infection, although our results in DF1 cells did not show evidence of virus replication. One possible explanation for the discrepancy in animal studies and *in vitro* studies is that the ACE2 protein in swine is not efficiently expressed in the respiratory tract, which is the most likely route of exposure, and the virus cannot efficiently attach and infect the exposed pig (54). Although results in swine are discordant, our studies using an avian cell line correlates closer with the swine challenge studies as we did not measure any virus in the cell supernatant that would be evidence of the virus completing the replication cycle, despite the possibility that the virus could attach to the modified cell line based on these previous studies.

The results with the horse ACE2 and TMPRSS2 genes showed no evidence of infection despite the relatively high sequence conservation of the horse ACE2 protein to human ACE2 at over 86%, which is higher than cats and Golden hamsters. Although the sequence similarity of human and horse ACE2 is high, the difficulty in challenging horses in a Biosafety level 3 animal facility has likely prevented the research from being performed. Our results provide additional support that horses are not susceptible to infection and do not need to be experimentally challenged.

Bats have been identified as likely reservoirs of both SARS-CoV-1 and MERS-CoV to humans through intermediate hosts including civet cats and dromedary camels, respectively (34–36). The SC2 virus appears capable to bind to Little Brown bat and Great Roundleaf Bat ACE2 as observed by positive immunostaining and transient virus replication. However, the TMPRSS2 protease found in these species may not be functional as it lacks the 5’ terminus found in human and other animals, including other bat species. Analysis of the GenBank record suggests that only partial sequence is available and that the gene was not properly annotated and thus the gene sequences used in these studies may not represent the true open reading frame. Further research is required to determine whether the anomaly is a sequence artifact. However, at least one report predicts low level fusion from Little Brown bat ACE2 compared to human ACE2, similar to the results described here (2). Further research is also underway to determine the contribution of different bat species ACE2 and TMPRSS2 as a barrier to SC2 infection.

As noted, SC2 appears to have a broad host range among mammals, however the full host range is unknown. Predictive *in silico* studies based on ACE2 analysis have described potential broad host tropism of the virus to numerous species including cat, goat and hamster (1, 2, 48, 49). These studies also predict many aquatic species including whales and dolphins to have high likelihood of binding to SC2 spike protein. *In silico* analysis of the TMPRSS2 protein is less predictive, but the protease activation of the SC2 spike protein is necessary for replication to occur. As noted earlier, proteases other than TMPRSS2 have been demonstrated to have the ability to cleave the spike protein. *In vivo* testing of many large domestic animals and wild animal species would be difficult, if not impossible, because of the requirement for work in a secure biocontainment facility. Therefore we propose this model could be utilized to screen many species for susceptibility to SC2 infection. Understanding the host range of SC2 is crucial to understanding the ecology of the virus and the role different species may play as reservoirs or bridge-species into humans. Species that can be infected also may be affected by disease. Our *in vitro* testing in DF1 ++ cells positively correlated with available *in vivo* challenge data. Taken together, the integration and expression of the ACE2 and TMPRRS2 from a target species in the otherwise non-permissive avian cell line provides a rapid and economical method to screen species for susceptibility to SC2.

## Acknowledgements

We thank Linda Moon, Scott Lee, Suzanne DeBlois, and Tim Olivier for excellent technical assistance. This research was supported by funding from USDA, ARS, CRIS project #6040-32000-066-00D.

The following reagent was deposited by the Centers for Disease Control and Prevention and obtained through BEI Resources, NIAID, NIH: SARS-Related Coronavirus 2, Isolate USA-WA1/2020, NR-52281. Vero African Green Monkey Kidney Cells (ATCC® CCL-81™), FR-243, was obtained through the International Reagent Resource, Influenza Division, WHO Collaborating Center for Surveillance, Epidemiology and Control of Influenza, Centers for Disease Control and Prevention, Atlanta, GA, USA.

